# HiCluster: A Robust Single-Cell Hi-C Clustering Method Based on Convolution and Random Walk

**DOI:** 10.1101/506717

**Authors:** Jingtian Zhou, Jianzhu Ma, Yusi Chen, Chuankai Cheng, Bokan Bao, Jian Peng, Terrence J. Sejnowski, Jesse R. Dixon, Joseph R. Ecker

## Abstract

3D genome structure plays a pivotal role in gene regulation and cellular function. Single-cell analysis of genome architecture has been achieved using imaging and chromatin conformation capture methods such as Hi-C. To study variation in chromosome structure between different cell types, computational approaches are needed that can utilize sparse and heterogeneous single-cell Hi-C data. However, few methods exist that are able to accurately and efficiently cluster such data into constituent cell types. Here, we describe HiCluster, a single-cell clustering algorithm for Hi-C contact matrices that is based on imputations using linear convolution and random walk. Using both simulated and real data as benchmarks, HiCluster significantly improves clustering accuracy when applied to low coverage Hi-C datasets compared to existing methods. After imputation by HiCluster, structures similar to topologically associating domains (TADs) could be identified within single cells, and their consensus boundaries among cells were enriched at the TAD boundaries observed in bulk samples. In summary, HiCluster facilitates visualization and comparison of single-cell 3D genomes.

**Availability:** https://github.com/zhoujt1994/HiCluster.git

## Introduction

In recent years there has been a rapid increase in the development of single-cell transcriptomic and epigenomic assays^1^, including single-cell/nucleus RNA-seq^2^, ATAC-seq^3,4^, bisulfite sequencing^5^ and Hi-C^6–11^. Such powerful techniques allow the study of unique patterns of molecular features that distinguish each cell type. Computational methods have been developed to identify different cell types in heterogeneous cell populations based on various molecular features such as transcriptome^12,13^, methylome^14^ and open chromatin^15–17^. However, unbiased and efficient algorithms for single cell clustering based on 3D chromosome structures are limited. In previous studies, principal component analysis (PCA) performed on both intra- and inter-chromosomal reads was unable to completely distinguish between four cancer cell lines^7^. An embedding method for single-cell Hi-C data has been specifically designed for capturing structural dynamics, allowing determination of the cell cycle state^18^. However, these data are continuous in nature and the approach has not explicitly been tested for the purpose of cell type identification.

Clustering of single cells based on Hi-C data faces three main challenges: (1) **Intrinsic variability**. 3D chromosome structures are highly spatially and temporally dynamic. Imaging-based technologies have suggested a large degree of heterogeneity of chromosome positioning and spatial distances between loci even within a population of the same cell type^19–22^. How this fluctuation between cells of the same cell type compares to fluctuations between different cell types remains unclear. (2) **Data sparsity**. The sparsity of single-cell Hi-C data is higher than most other types of single-cell data. State-of-the-art single-cell DNA assays typically cover only 5-10% of the linear genome. Since Hi-C data is represented as two-dimensional contact matrices, this level of sensitivity leads to coverage of only 0.25-1% of all contacts to be captured. (3) **Coverage Heterogeneity**. It is often observed that the genome coverage of cells extends over a wide range within a single cell Hi-C experiment. We find this bias often acts as the leading factor to drive clustering results, making it difficult to systematically eliminate. For example, this bias could be alleviated by removing the first principal component (PC1) before clustering and visualization. However, PC1 is not guaranteed to represent only cell coverage in these experiments as it may also contain information related to other biological variables.

To address these challenges, we developed a new computational framework, HiCluster, to cluster single-cell Hi-C contact matrices. To overcome the sparsity problem, we performed two steps of imputation on the chromosome contact matrices to better capture the topological structures. To solve the heterogeneity problem, we selected only the top-ranked interactions after imputation, which were proved to be sufficient to represent the underlying data structure. This framework significantly improved upon the clustering performance using low coverage datasets as well as facilitated the visualization and comparison of chromosome interactions among single cells.

## Results

### Overview of HiCluster

As shown in **Fig. 1**, HiCluster consists of four major steps. In the first step, every element of the contact matrix is replaced by the weighted average of itself and its surrounding elements, in a type of linear convolution. Then a random walk (with restart) algorithm is applied to smooth the signal to further capture both the local and global information of the contact maps. In particular, the convolution step only allows the information to pass among the linear genome neighbors, while the subsequent random walk step aides information sharing among the network neighbors. To alleviate the bias introduced by uneven sequence coverage, we only keep the top 20% interactions after the imputation. Finally, we project the processed contact matrices onto a shared low dimensional space, so that the topological structure of the 3D chromosome contacts can be compared between cells and used for further clustering and visualization.

**Figure 1.**
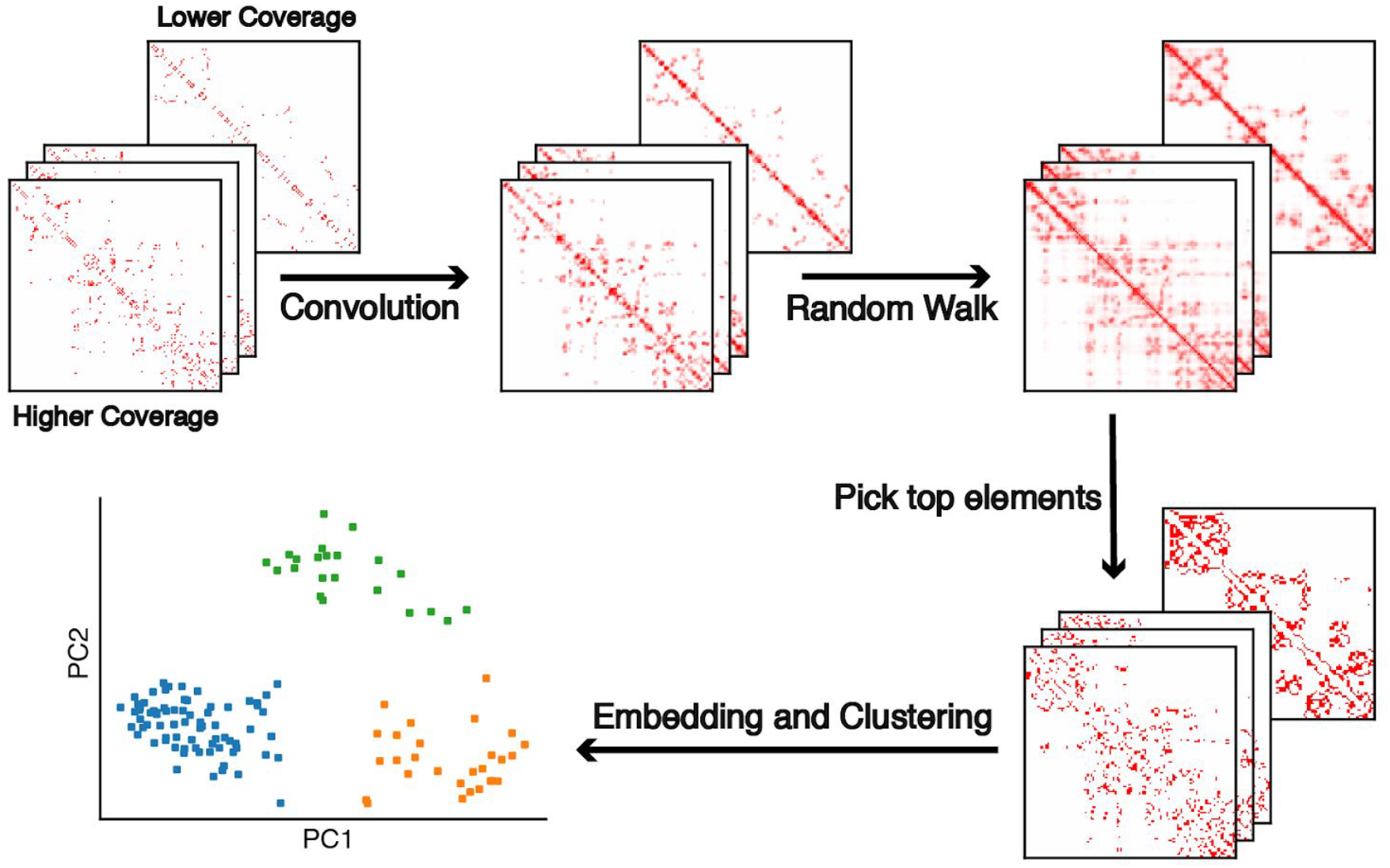
The workflow of HiCluster. The contact matrices of each single cell are smoothed by two steps of imputation that include convolution and random walk; these are based on the neighboring bins of a linear genome and long-range connections, respectively. To alleviate the coverage bias, only top 20% elements of the imputed matrices are selected. Each single cell matrices is then projected into the same space and then clustering is performed to identify distinct cell types.

### HiCluster improves clustering performance on simulated data

To explore the combinatorial effects of different levels of coverage and resolution, we first applied our algorithm to a set of simulated single-cell Hi-C data. We noticed that direct sampling from the Hi-C contact matrices of bulk cells leads to a relatively lower sparsity and heterogeneity (**Fig. S1**), which often yields more accurate clustering results compared to real single-cell data. The real data concentrated more on specific loci in each cell, and the individual loci were different between different cells (**Fig. S1A**). On the contrary, the simulated cells from bulk data often had more evenly distributed contacts (**Fig. S1B**). Therefore, we controlled the sparsity of each simulated contact matrix and added noise to the contact-distance curves to better mimic the sparsity and noise of real data (**Methods**). As shown in **Fig. S1G**, when considering the first two principal components (PCs), the simulated cells generated were indistinguishable from real single cells of the same cell type.

In our simulation we performed downsampling from bulk Hi-C experimental data from two studies. Rao et al. 2014^23^ examined seven human cell types (GM12878, IMR90, HMEC, NHEK, K562, HUVEC and KBM7) while Bonev et al. 2017^24^ examined three mouse cell types (embryonic stem cells (ESC), neural progenitor cells (NPC) and cortical neurons (CN)). We downsampled each dataset to 500k, 250k, 100k, 50k, 25k, 10k, 5k contacts respectively, and used 1 Mbp and 200 kbp resolution contact maps to test our algorithm. At each coverage level and resolution, we generated 30 simulated cells for each cell type. We evaluated the ability of HiCluster compared with PCA to recover the correct cell type in an unsupervised way. The adjusted rand index (ARI) was used to measure the accuracy of clustering. As shown in **Fig. 2** and **Fig. S3**, in both datasets, HiCluster consistently performed better than PCA. The performances began to be impaired with fewer than 25k contacts, and a complete loss of clustering ability is observed at 5k contacts. We also found that 1 Mbp resolution performed better than 200kbp (**Figs. S4C, D**), suggesting that lower sparsity (lower resolution) may be sufficient to distinguish cell types. Thus, we used 1 Mbp resolution in all subsequent experiments.

**Figure 2.**
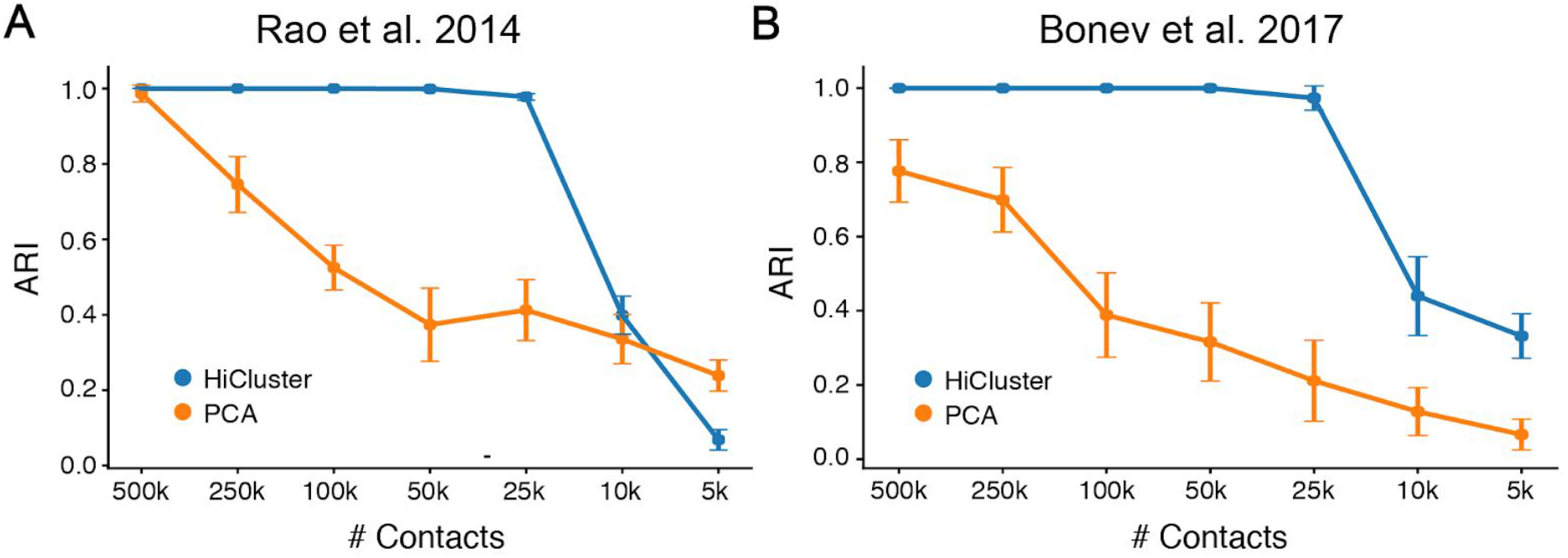
The performance of HiCluster and PCA on simulated data. Bulk Hi-C contact matrices in Rao et al. 2014 (A) and Bonev et al. 2017 (B) were sampled to 100k, 50k, 25k, 10k and 5k contacts respectively. The clustering performance is measured by Adjusted Rand Index (ARI).

### HiCluster has superior performance on published single-cell Hi-C data

Next, we evaluated our analysis framework using authentic single-cell Hi-C datasets. Thus far there have been three published studies focusing on single-cell chromosome structures with analyses of multiple cell types. Ramani et al.^7^ used a combinatorial indexing protocol to generate single-cell Hi-C libraries from thousands of cells for four human cell lines (HeLa, HAP1, GM12878 and K562). The number of contacts captured in each cell ranged from 5.2k to 102.7k (median 10.0k). Flyamer et al.^10^ performed whole genome amplification after ligation and detected 6.6k to 1.1m contacts per cell (median 97.3k) in mouse zygotes and oocytes. Tan et al.^11^ developed an optimized protocol also using whole genome amplification and obtained data with a median coverage of 513.0k contacts. Since the last benchmark dataset (Tan) had relatively high coverage, either simple PCA (**Fig. S5**) or chromosome compartment score^11^ easily allowed cell types to be distinguished. Due to cost considerations, it is still challenging to achieve such depth of genome coverage. Therefore, we focused on the first two datasets with lower coverage (Ramani and Flyamer) to test the utility of our computational framework.

We compared our algorithm with three baseline methods: PCA, HiCRep+MDS and the Eigenvector method. Liu and colleagues compared different tools to calculate the similarity of single cell Hi-C matrices and concluded that HiCRep followed by multidimensional scaling (MDS) performed the best among all the single-cell embedding methods they investigated^18^. The chromosome compartments are also considered to be cell type specific based on the bulk Hi-C experiments, and the first eigenvector of contact matrix is widely used to represent these compartment features^23,25,26^. HiCluster outperformed the baseline methods on both datasets in terms of better visualization (**Figs. 3A, B**) and improved ARI (**Figs. 3C, D**). In the mouse dataset (Flyamer), HiCluster made a significant distinction among all three cell types (**Fig. 3A**); while in the human dataset (Ramani), the algorithm separated K562 and HAP1 better in the first two PC dimensions (**Fig. 3B**). It is also worth commenting on the scalability of each methods. Computing the HiCRep similarity required 8 hours (Flyamer) and 4.5 days (Ramani); however, the HiCluster and other methods consumed around 30 seconds (Flyamer) and 60 seconds (Ramani) (**Fig. S6**). Additionally, we carried out the same experiments on each chromosome separately and noticed that almost every chromosome showed advanced separation on the mouse dataset (**Fig. 3E**), while only one chromosome showed significant improvement on the human dataset (**Fig. 3F**). These results may suggest that to separate cells using global chromosome structure differences (e.g., oocytes and zygotes), the information provided by a single chromosome might be sufficient, but to distinguish more complex cell types, a combination of different chromosomes or a more careful feature selection is necessary.

**Figure 3.**
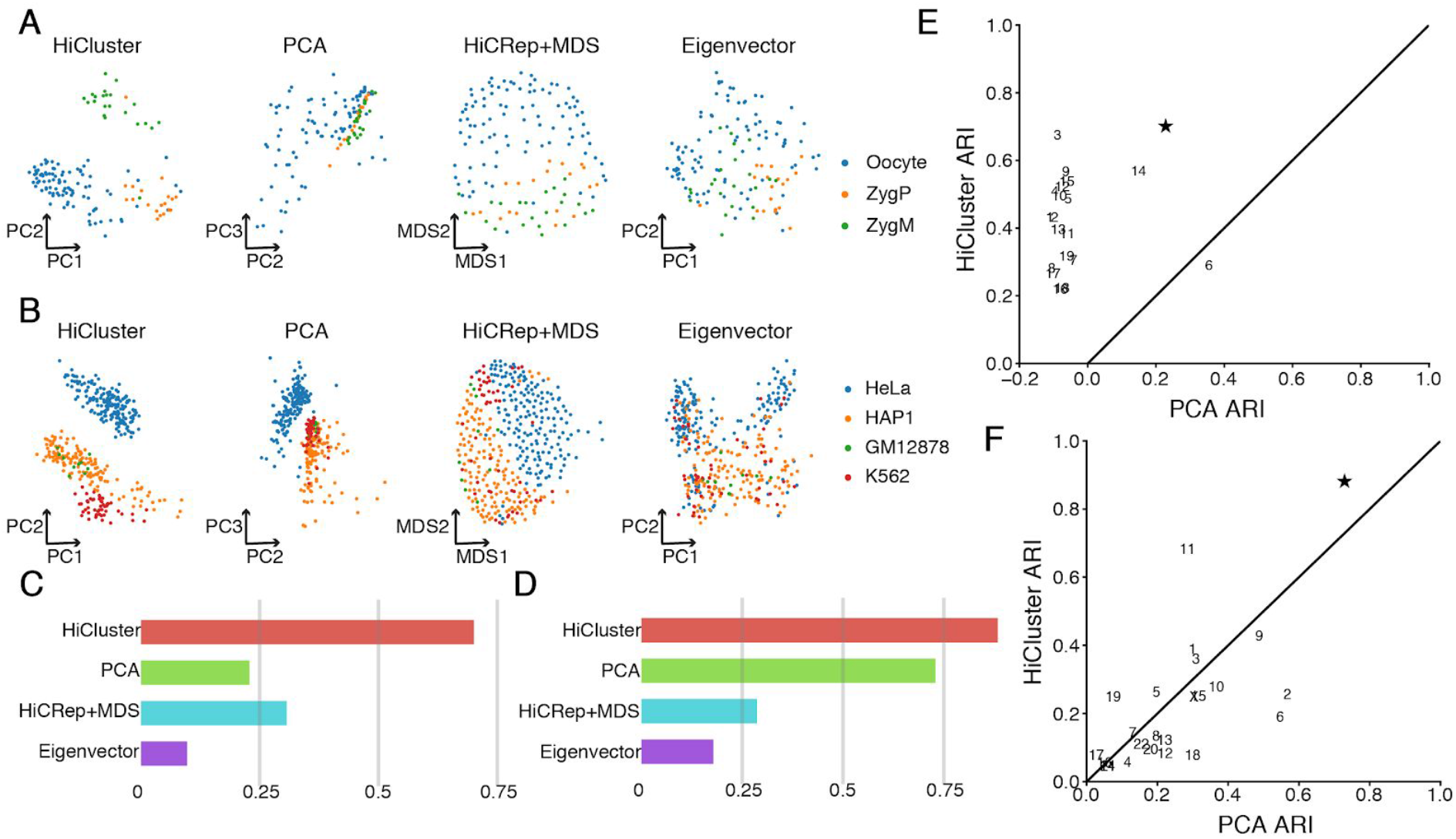
The performance of HiCluster and baseline methods on real single-cell Hi-C data. For Flyamer et al. 2017 (A, C, E) and Ramani et al. 2017 (B, D, F), the embedding (A, B) and ARI of clustering (C, D) are shown. For HiCluster and Eigenvector, the embeddings are shown in PC1 and PC2 space. For PCA, the embedding is shown in PC2 and PC3 space. (E, F) The performance of HiCluster and PCA on each chromosome (indicated by chr. numbers). The ARI using all chromosomes (indicated by star symbol).

We also visualized the weights of each element in the contact matrices when computing the final PCs (whitening matrices). In general, the weights for PC1 were uniformly distributed parallel to the diagonal (**Fig. S7A**), which suggested it captures the information of the contact-distance curve and might correspond to the variance resulting from cell-cycle or other relevant biological effects^9^. This is also corroborated by the observation that cells with greater PC1 values tended to have a higher frequency of short-range contacts, while smaller PC1 inclined to correspond to a higher frequency of long-range contacts (**Fig. S7B**). On the contrary, the weights for computing PC2 showed region specificity (**Fig. S7A**), which may indicate its correlation with compartment strength. These finding also explained why the oocytes and zygotes in Flyamer 2017 are dominantly separated by PC1, where the contact distance curve differ between cell types; meanwhile, PC2 performed better partition of the cancer cell lines in Ramani 2017.

Next, we wanted to evaluate the contribution of each step to the final clustering performance. For the three major steps of the pipeline, we tested all possible combinations of one or two steps of the three. More specifically, we compared our framework with PCA (with none of the steps), CONV (convolution only), RWR (random walk only), CONV_TOP (convolution and select top elements), RWR_TOP (random walk and select top elements) and CONV_RWR (convolution and random walk). From **Fig. 4**, we concluded that all three steps are necessary to achieve the current visualization (**Figs. 4A, B**) and clustering accuracy (**Figs. 4C, D**). The necessity of these steps is more evident when using the mouse dataset. Notably, for the methods with skipped steps, we chose the best ARI among a wide range of PC dimensions and clustering methods; while for the framework with three steps, we fixed the clustering part to be K-Means on 10 PCs.

**Figure 4.**
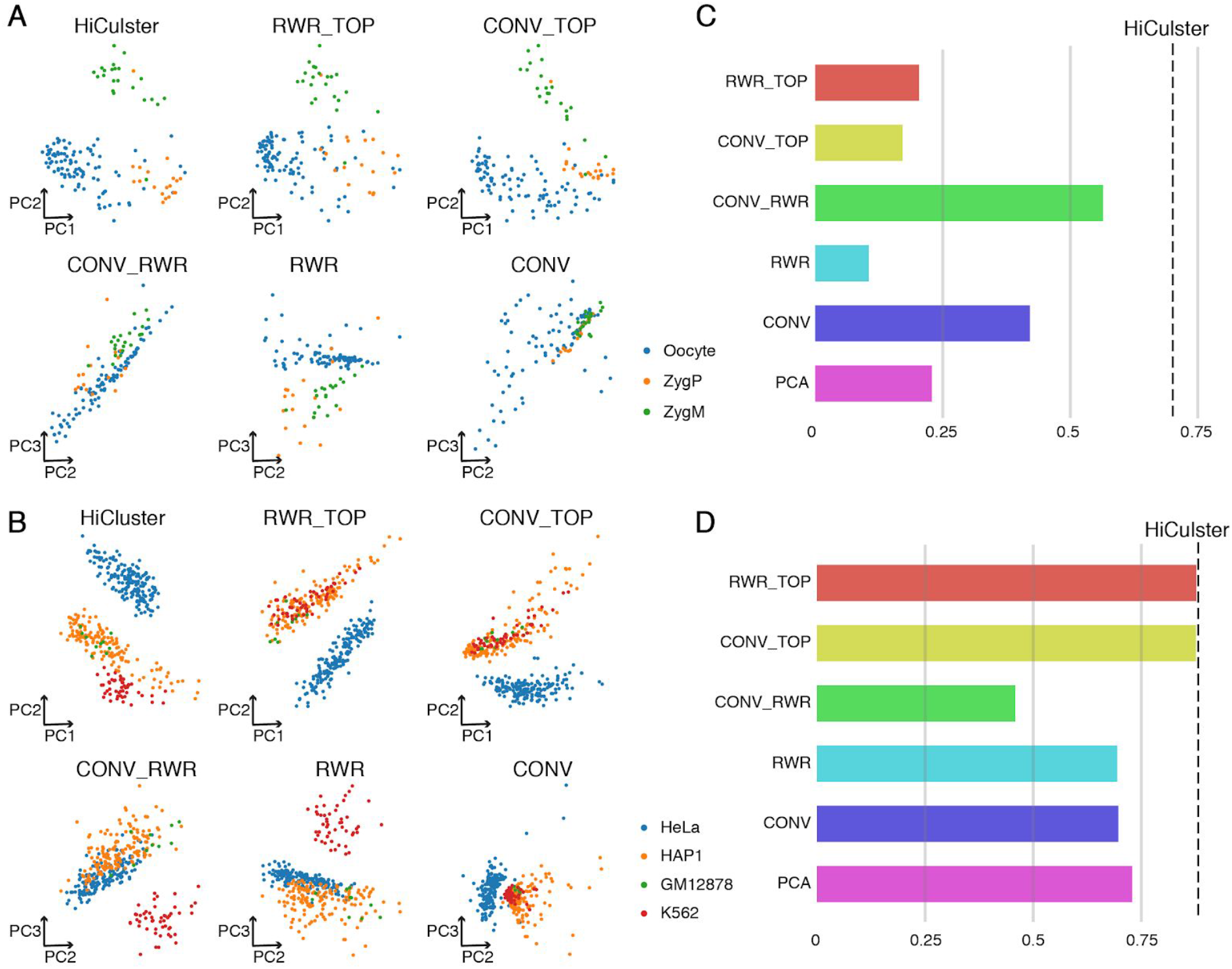
The performance of HiCluster applying different combinations of substeps. For Flyamer et al. 2017 (A, C) and Ramani et al. 2017 (B, D), the embedding (A, B) and ARI of clustering (C, D) with each combination of steps are shown. (A, B) The embeddings of combinations with TOP step are shown in PC1 and PC2 spaces, without are shown in PC2 and PC3 spaces. (C, D) The black dash line denotes the performance with all steps.

### HiCluster allows visualization of structural difference in single cells

The most popular method to interpret and validate identified cell clusters in single cell experiments is to analyze known marker genes. Gene expression is directly measured in single-cell RNA-seq data and promoter, gene body ATAC-seq signals or cytosine methylation ratios can also be used to infer the cluster-specific genes in single-cell open chromatin and methylome data. Similarly, in single-cell Hi-C data, the differential chromosome interactions could serve as cell-type markers. With the single-cell Hi-C data, imputed contact matrices from every single cluster can be merged, where we observed square patterns that are visually similar to the topologically associating domains (TADs) identified in bulk Hi-C experiments along the diagonal. However, since the existence of TAD remains unclear in single cells, and accurate identification of the structures were limited by data sparsity, we referred to this featured pattern as TAD-like structures (TLSs) hereafter. Thus, differential TLSs could be applied to characterize different cell types. For instance, as demonstrated in **Fig. 5** and **Fig. S8**, a TLS at chr9:133.6M-134.2M is observed in 9 of 10 K562 cells but in two of the GM12878 cells. This structure difference is concordant with the bulk Hi-C data from the same cell lines. Gene expression and H3K4me1 signals which marks active enhancers are also higher in K562 within this TLS.

**Figure 5.**
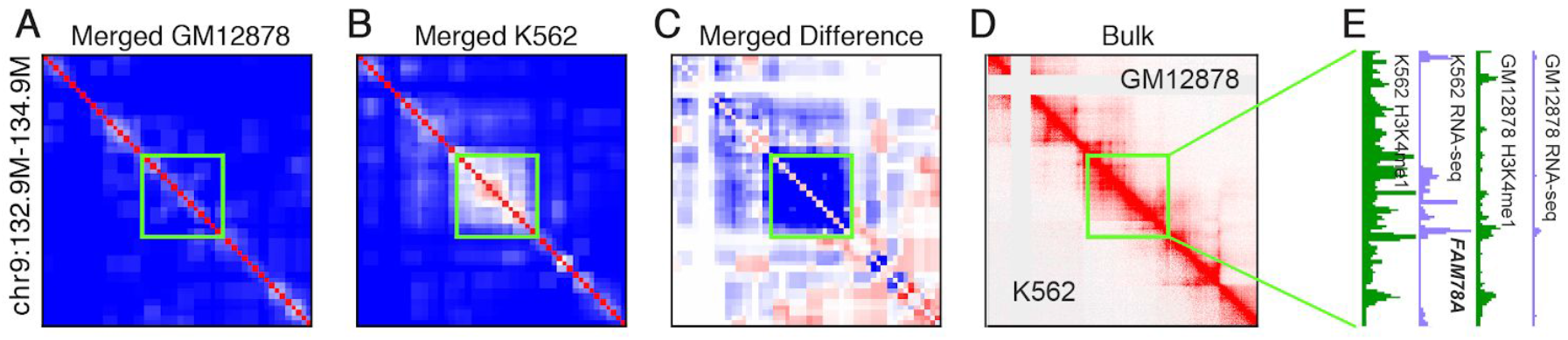
Visualization of contact matrices at chr9:132.9M-134.9M. Merged contact matrices from 10 GM12878 (A) and K562 (B) cells after imputation. (C) The difference between (A) and (B). (D) The contact matrices of bulk cell lines. (E) The RNA-seq and H3K4me1 signals in both cell lines.

Structural differences are also observed near differentially expressed genes between the two cell types, including *CXCR4* and *ZBTB11. CXCR4* is a chemokine receptor that enhances cell adhesion, which is highly expressed in non-cancer cells (GM12878) comparing to cancer cells (K562)^27^. With HiCluster imputation, a TLS surrounding *CXCR4* was detected in 6 of 10 GM12878 cells but only 2 of 10 K562 cells (**Fig. S9, Methods**). Intriguingly, an H3K4me1 peak was detected in bulk GM12878 but not K562 at the other boundary of the TLS, which may indicate the potential interaction between the gene and its enhancer. Similarly, a TLS whose boundary located at *ZBTB11* was observed in more GM12878 cells than K562 cells (**Fig. S10**). Consistently, more H3K4me1 peaks within this TLS were also detected in the bulk GM12878 sample.

Next, we examined if the imputation based on HiCluster could facilitate the systematic identification of TLSs in both simulated and real single cells. We first leveraged Bonev et al. data for bulk ESC and NPC, and downsampled them to 1 Mbp, 500 kbp, 250 kbp, 100 kbp contacts per cell. We applied HiCluster on contact matrices and run TopDom^28^ to detect TLSs in every single cell. A TAD in NPC that splits into two TADs in ESC was selected to test the performance of TLS-calling (**Fig. 6A**). The visualization of single cell TLSs was significantly improved after HiCluster smoothing (**Fig. 6B**), and the alternative boundary was captured in more cells (**Fig. 6C**). Next, we applied HiCluster to analyze single-cell Hi-C data from Nagano et al., which contains 1,992 mouse ESCs across different stages of cell-cycle^9^. The dataset was sequenced with high coverage and enabled us to statistically analysis the dynamic of TADs location within single cells. We identified TLSs on the contact matrices smoothed by HiCluster at 40 kbp resolution with TopDom, and for each bin, we counted the number of cells in which the bin was determined as a TLS boundary. We observed non-zero probability for almost all bins to be a TLS boundary in single cells, and these probabilities peaked at the Ctcf and cohesin binding sites, and the TAD boundaries described in bulk Hi-C (**Fig. 6D**), which is in agreement with the conclusions of a recent imaging study^29^. This signal was significantly enhanced after convolution and random walk (**Fig. 6D**), which further highlighted the potential application of HiCluster to study single cell chromosome structure.

**Figure 6.**
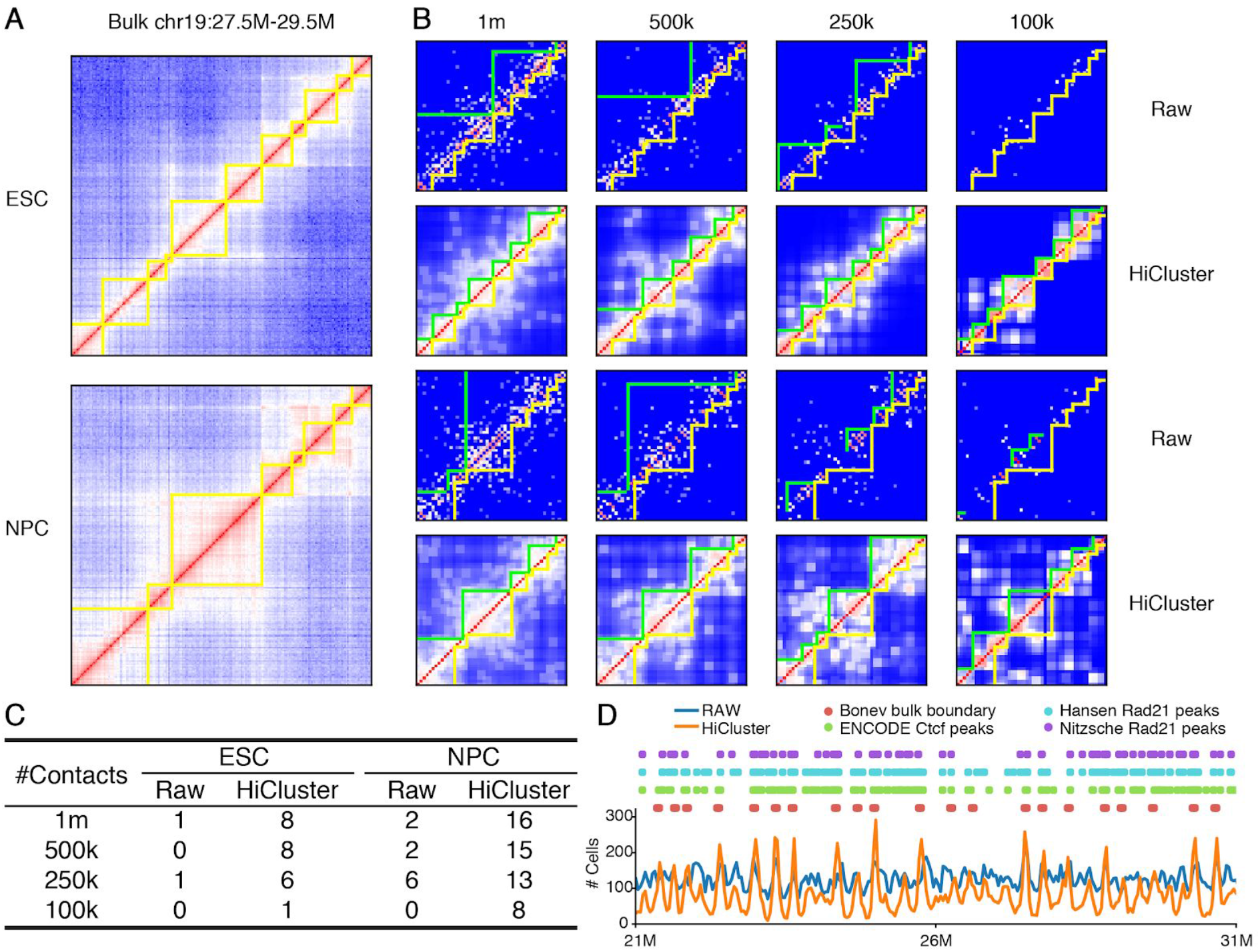
HiCluster facilitates identification of domain-like structures (TLSs). (A) The contact matrices at chr19:27.5M-29.5M of bulk Hi-C data with alternative TADs in ESC and NPC. (B) The downsampled contact matrices with 1 Mbp, 500 kbp, 250 kbp, 100 kbp total contacts per cell before and after HiCluster imputation. The green lines indicate the TLSs called from the plotted matrix, and yellow lines represent the TADs called from bulk data of the corresponding cell type. (C) The number of downsampled ESCs with TLSs at chr19:28170000-28530000 and chr19:28530000-28770000, or downsampled NPCs with TLSs at chr19:28170000-28770000 being identified before and after HiCluster imputation. (D) The number of single ESCs (1007 in total) with TLSs boundary identified at each genome bin are shown by lines. The position of Ctcf ChIP-Seq peaks, Rad21 ChIP-Seq peaks and TADs boundaries identified in bulk data are presented by dots.

Our imputation method also helps visualize the signature of chromatin structures within specific cell type. *Sox2* is a classic marker gene of ESCs, and the chromosome structure around this gene is unique to ESCs^24^. Specifically, *Sox2* is located within a large TAD in NPC (**Fig. S11B**), which is split into two smaller TADs in ESCs, with *Sox2* located at the boundary between the two smaller TADs (**Fig. S11A**). Stevens et al.^8^ carried out Hi-C analysis of eight single haploid mouse ESCs. A median of 49.4 kbp long-range intra-chromosome contacts was detected (21.0k to 78.0k). Although this study provided superior coverage among the current single-cell Hi-C experiment, the limited number of cells examined made it difficult to observe the interaction pattern surrounding the *Sox2* even if contact matrices from all cells are merged (**Fig. S11C**). However, after the imputation using the HiCluster framework, the TLS boundaries at *Sox2* are observed in 4 of the 8 cells (**Fig. S11E**). Merging the imputed matrices reveals the known domain splitting pattern near *Sox2* (**Fig. S11D**). A similar interaction pattern is also observed near another ESC marker *Zfp42* (**Fig. S12**).

## Discussion

To advance our understanding of the role of genome structure in cell-type specific gene regulation, new computational tools are needed for deconvolution of single cell Hi-C data. We describe a novel computational approach for cell type clustering, HiCluster, that requires only sparse single cell Hi-C contact data. In the HiCluster framework, the chromosome interactions are considered as a network. The contact information is first averaged in the linear genome. A random walk is then used to propagate the smoothed interaction throughout the graph and further reduce the sparsity of the single cell contact matrices. HiCluster performed significantly better than existing methods in clustering single-cell data into constituent cell types and facilitated identification of local chromosome interaction domains.

A major challenge in clustering single cell Hi-C data is the sparsity of the contact matrices. To overcome this limitation, HiCluster uses both a linear smoothing and a random walk step. Similar methods have been utilized for smoothing bulk Hi-C data, including HiCRep which took the average of genome neighbors before computing the correlation of two Hi-C matrices^30^, and GenomeDISCO which provided a network representation of Hi-C matrices and used random walk to smooth it^31^. Liu et al.^18^ systematically evaluated these methods for single cell Hi-C data embedding. However, since they used a cell similarity matrix that is embedded by MDS, the data are generally continuous under their low-dimensional representation and are unable to present explicit clusters for each cell type. Our HiCluster framework combines the advantage of both HiCRep and GenomeDISCO and provides a flexible pipeline to resolve the clustering of Hi-C data, where some components (e.g. embedding) can be further tuned and improved when the algorithm is applied to more specific and challenging situations such as tissues with greater cell type complexity.

Published single cell Hi-C datasets have employed cell lines which contain relatively large 3D genomic structural differences, simplifying the cell clustering problem. In practice, heterogeneous tissues with more closely related cell types, such as brain tissue, might pose a much greater challenge than cell lines. For cell clustering using complex tissues, further improvement of clustering algorithms and feature selection would be necessary. An alternative approach would be to simultaneously profile 3D genome architecture along with other “omic” information in the same cell, such as jointly profiling chromatin conformation and DNA methylation^32^. While such single cell multi-omic data modalities may provide the information content necessary to deconvolute cell types while preserving 3D structural information^33^, they can also be more costly to perform, and more technically challenging to carryout.

We noted that the smoothing and random walk steps aid in visualization of chromosome contact maps in single cells. Such visualization can facilitate analysis of the variability in features of 3D genome organization between cells. Previous studies using bulk cell lines have reported the existence of several 3D structural features: megabase level A/B compartments, sub-megabase level TADs, and kilobase level loops^23,25,34,35^. In our study, visualizing the smoothened HiCluster results revealed the existence of TLSs in specific cells. The boundaries of these structures were variable between cells. However, the boundaries shared between TLSs in individual cells corresponded to TAD boundaries identified in bulk Hi-C studies. These results would support recent imaging studies^29^ which suggested that TLSs exist in single cells, and their boundaries in individual cells are variable but non-random.

## Methods

### Data processing

For Ramani 2017, interaction pairs and cell quality files of combinatorial single cell Hi-C library ML1 and ML3 were downloaded from GSE84920. Interaction pairs for Flyamer 2017, Stevens 2017 and Tan 2018 were downloaded from GSE80006, GSE80280 and GSE117876, respectively. Interaction pairs for diploid ESC cultured with 2i in Nagano 2017 were accessed from https://bitbucket.org/tanaylab/schic2/src/default/. Given a chromosome of length *L* and a resolution *r*, the chromosome is divided into 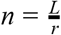 non-overlapping bins. Hi-C data is represented as a *n* × *n* contact matrix *A*, where *A_ij_* denotes the number of read-pairs supporting the interaction between the *i*-th and *j*-th bins of the genome. For each dataset, contact matrices were generated at 40kbp and 1 Mbp resolutions for each chromosome and each cell. Total contacts of the cell were counted as the non-diagonal interaction pairs in intra-chromosomal matrices. As quality control, we ruled out the cells with less than 5k contacts. Also, for a single chromosome whose length is *x* Mb, we required the number of contacts to be greater than *x*, in order to avoid the chromosomes with too few contacts. We only kept cells where all chromosomes satisfied this criterion.

### Simulations

We downsampled bulk Hi-C data to simulate datasets with similar sparsity and heterogeneity of single cells. Bulk MAPQ30 contact matrices were extracted from Juicerbox at 100kbp resolution for datasets Rao et al. 2014 and Bonev et al. 2017, respectively. Contact matrices for each cell type at 200kbp and 1 Mbp resolution were calculated by merged bins in the 100kbp-resolution matrices.

#### Normalization

SQRTVC normalization was applied to the bulk contact matrices to deal with the coverage bias along the genome. The normalized contact matrices *B* are computed by

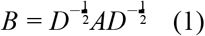

where *D* is a diagonal matrix where each elements *D_ii_* is the sum of the *i*-th row of *A*.

#### Sparsity controlling

After directly downsampling the read counts from *B*, we still observed a significantly lower sparsity of simulated data comparing to real single-cell data with the same number of contacts in total (**Figs. S1A, B**). Therefore, we further controlled the sparsity during sampling to make the simulated data more similar to the real data. Leveraging Ramani et al. 2017 and Flyamer et al. 2017 datasets, we fit a linear relationship between total contacts *C* and sparsity *S* at log scale (**Fig. S2**),

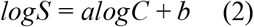

To generate a simulated dataset with the median contact counts to be *M*, for each simulated single cell we uniformly sampled *t* from *logM* – 0.5 to *logM* + 0.5 and set the total contacts number of the cell as *C* = *e^t^*. The sparsity of the cell *S* was computed based on (2). The sampled new contacts are randomly assigned to different chromosomes based on the contact numbers of each chromosome in a particular cell type in the bulk cell dataset.

#### Adding random noise

After controlling the sparsity of the contact matrices, the heterogeneity of the simulated cells is still not high enough to mimic the complex situations within the real data. To address the problem, we add noise to the contact frequency through Contact-Distance Curve, which describes the values in the contact matrices changed with respect to their distance to the diagonal. More specifically, we generated a random vector *R* of length *n*, where *n* is the bin number of the contact matrix. The values in *R* ranges from −*k* to *k* following a uniform distribution, where *k* denotes the noise level. Then, the normalized bulk contact matrix *B* was rescaled linearly to the noisy representation *E* by *E_ij_* = *B_ij_* × *R_|j−i|_*. Finally, based on *E*, we sampled *S* positions to be non-zero candidates based on **Eq. 2**, and distributed the *C* simulated contacts to these positions.

### HiCluster

#### Convolution-based imputation

Imputation techniques are widely adopted in single-cell RNA-seq data to improve the data quality based on the structure of the data itself. For HiCluster, the first step is to integrate the interaction information from the genomic neighbors to impute the interaction at each position. The missing value in the contact matrix could be due to experimental limitations of material dropout, rather than no interactions. Since the genome is linearly connected, our hypothesis is that the interaction partners of one bin may also be close to its neighboring bins. Thus, we used a convolution step to inference these missing values. Specifically, given a window size of *w*, we applied a filter *F* of size *m* × *m*, where *m* = 2*w* + 1, to scan the contact matrix *A* of size *n* × *n*. The elements in the imputed matrix *B* is computed by

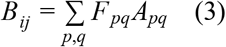

where *i* − *w* ≤ *p* ≤ *i* + *w, j* − *w* ≤ *q* ≤ *j* + *w*. In this work, all the filters are set to be all-one matrices, which is equivalent to taking the average of the genomic neighbors. However, the filters could be tuned to incorporate different weights for elements during imputation. For instance, the elements located further from the imputed elements could be assigned smaller weights. The window size *w* was set to 1 for 1 Mbp resolution maps.

#### Random-walk-based imputation

Random walk with restarts (RWR) is widely used to capture the topological structure of a network^36^. The random walk process helps to infer the global structure of the network and the restart step provides the information of local network structures. What Hi-C data fundamentally describe is the relationship between two genomic bins, which can be considered as a network where nodes are the genomic bins and edges are their interactions. Different from the convolution step which taking information from the neighbor on the linear genome, the random walk step considers the signal from the neighbor with experimentally measured interactions. The imputed matrix *B* defined in Eq (3) is first normalized by its row sum.

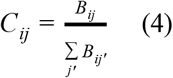

We use *Q_t_* to represent the matrix after the *t*-th iteration of random walk and restart. Then the random walk starts from the identity matrix *Q*_0_ = *I*, and *Q_t_* is computed recursively by

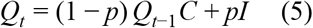

where *p* is a scalar representing the restart probability to balance the information between global and local network structures. The random walk with restart was performed until ║*Q_t_* − *Q*_*t*−1_║_2_ ≤ 10^−6^. Each element *Q_ij_* in the matrix after convergence signifies the probability of random walk to reach the *j*-th node when starting from the *i*-th node.

#### Embedding

Since the coverage of the matrices from each cell is different, the sparsity and scales of the matrices after random walk is also distinct. Thus, after random walk, a threshold *t* was chosen to convert the real matrix *Q* into binary matrix *Q_b_*. The threshold *t* was set to be the 80th percentile of *Q* for all the analysis, and its impact is discussed in **Fig. S4**. This is a crucial step since it facilitates us to choose the most conserved and reliable interactions in each cell. Then the *n* × *n* matrix *Q_b_* is reshaped to 1 × *n*^2^ and the matrices from *m* different cells were concatenated into a *m* × *n*^2^ matrix. In the last step, PCA was used for projecting the matrix into a low dimensional space and produce the embedding of the cells. Each single chromosome was embedded separately and the embedding of all chromosomes was concatenated at last and another PCA was applied to derive the final embedding. The whitening matrices for the two steps of PCA were multiplied, and the dot product representing the weight of each element in the contact matrices for computing each PC was visualized.

### Baseline methods

#### PCA

The raw contact matrices of each cell were log2 transformed and reshaped to 1 × *n*^2^. The matrices from *m* different cells were concatenated into a *m* × *n*^2^ matrix. The matrix for each chromosome was PCA transformed and concatenated at last, and another PCA was applied to derive the final embedding with all chromosomes.

#### HiCRep+MDS

HiCRep 1.6.0 was installed from bioconductor. For each chromosome, the raw contact matrix of each cell were log2 transformed and smoothed with a window size of 1. The stratum adjusted correlation coefficient (SCC) were computed between each pair of smoothed matrices. The median of SCC distances across all chromosomes were transformed to euclidean distances by Eq (6).

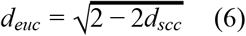

The Euclidean distance matrix was then embedded into two dimensions with MDS.

#### Eigenvector

The raw contact matrices of each cell were log2 transformed. PCA was performed on each cell and Z-score transformed PC1 was kept as features of the cell. We computed the mean CpG content of the bins with positive and negative features respectively and reversed the features if the negative features corresponded to higher CpG content. The features from *m* different cells were concatenated into a *m* × *n* matrix and PCA transformed.

### Identification of TLSs/TADs

All TLSs/TADs were computed by TopDom with a window size of 5. TADs in bulk ESC and NPC were identified at 10kbp resolution. All HiCluster imputations on contact matrices were performed at 40 kbp resolution with a window size of 1 and restart probability of 0.5, without selecting the top elements. The number of cells with differential TLS were counted as cells with contacts at greater than 40% elements in the green box. The number of TLSs in **fig. 6C** were counted as TLS identified with both boundaries within 0.25-fold of corresponding TAD boundaries identified in bulk data.

## Acknowledgments

We thank Dr. Jie Liu for specifying the details in his algorithm. J.R.E. is an investigator of the Howard Hughes Medical Institute.

